# The Longest 3D-preserved Plumage Reveals Stepwise Pennaceous Feather Evolution

**DOI:** 10.64898/2026.05.03.722549

**Authors:** Yanyun Zhang, Jiawei Tang, Natalia Jagielska, Shuo Wang

## Abstract

Powered flight is a critical innovation associated with the evolutionary transition from non-avialan theropods to birds, yet how early feathers gave rise to modern pennaceous feather structures with optimized aerodynamic performance remains unclear. Here we report a three-dimensionally preserved pennaceous feather from the Burmese amber (∼99 Ma) that exceeds 105 mm in preserved length, representing the longest known feather preserved in amber. It shows symmetrical vanes with densely packed barbs, indicating a derived pennaceous branching architecture, but lacks interlocking barbules and exhibits incompletely differentiated rachis and barbs, implying limited aerodynamic performance. This combination of advanced branching organization and incomplete tissue differentiation indicates asynchronous evolution of pennaceous feathers, in which branching organization, elongation and vane organization preceded the acquisition of interlocking barbules and fully differentiated cortical and medullary tissues required for aerodynamic function. These findings provide direct fossil evidence for stepwise, modular evolution of pennaceous feathers. They suggest that aerodynamic optimization of flight-related feather structures may not have been the primary driver of pennaceous feather branching.

**Significance statement:** How pennaceous feathers became mechanically specialized for powered flight remains poorly understood. Here we report that the longest known three-dimensionally preserved pennaceous feather from Burmese amber (∼99 Ma) exhibits a symmetrical but non-interlocking vane and limited tissue differentiation in the rachis and barbs, capturing a previously undocumented transitional stage in pennaceous feather evolution. This structural mosaic provides direct fossil evidence that aerodynamic performance was optimized through stepwise, modular evolution of pennaceous feather structures.

## Introduction

Feathers, the primary integumentary structures of non-avialan dinosaurs and birds, are highly diverse in morphology and function, with form strongly constraining functional performance [1]. The emergence of pennaceous feathers is widely regarded as a key evolutionary innovation associated with the origin of remiges and/or rectrices during the theropod–bird transition [2-5] and is therefore commonly regarded as a morphological proxy for flight capability [6, 7]. However, how early feathers gave rise to modern pennaceous feather structures with optimized aerodynamic performance remains unclear.

Large pennaceous feathers are known primarily from pennaraptoran theropods [8], whereas more basal taxa such as *Yutyrannus* and *Sinosauropteryx* preserve only filamentous integuments without differentiation of rachises and vanes, consistent with the absence of aerodynamic function [9, 10]. Most large fossil feathers are preserved as carbonized compressions [4, 11-13], in which fine structural details such as barbule hooklets and keratinous ultrastructure are seldom retained [14]. Accordingly, aerodynamic performance has largely been inferred from overall feather morphology, with limited direct assessment of how detailed feather architecture influences aerodynamic function [15].

Amber inclusions, by contrast, preserve feathers three-dimensionally and often retain keratinous tissues and microstructural features such as melanosomes with exceptional fidelity [16-20]. Amber-preserved feathers show that incompletely differentiated rachises and barbs were more widespread in early feathers than in extant forms [18, 19, 21]. However, large pennaceous feathers are rarely preserved in amber because of the limited size of most resin pieces, and most known inclusions are small downy or contour feathers only 5–20 mm long [16-20].

Here we describe the longest known three-dimensionally preserved pennaceous feather from the Burmese amber (Cenomanian, ∼99 Ma) [22]. Exceeding 105 mm in preserved length and retaining nearly three-quarters of its original extent, the specimen combines a symmetrical but non-interlocking vane with incompletely differentiated rachis and barbs. This structural mosaic reveals a combination of derived overall organization and primitive architecture, capturing a transitional stage in the assembly of pennaceous feathers and clarifying how aerodynamic function evolved in early theropods.

## Materials and Methods

### Feather sampling

The amber sample (ECNU A0033) was donated in 2016 by an anonymous amateur collector to the Biology Museum of East China Normal University (ECNU), Shanghai, China, along with other feather specimens previously documented [18]. The specimen is not association with the Myanmar Economic Corporation, a military-owned conglomerate, thereby ensuring full compliance with the moratorium issued by the Society of Vertebrate Paleontology [23].

### Microscopy and Measurements

Because of the extremely low contrast of the feather inclusion, high-resolution CT imaging was insufficient to resolve fine structural details. Morphological characterization was therefore conducted using light microscopy and scanning electron microscopy (SEM). Non-destructive macroscopic imaging was performed using a Nikon SMZ25 stereomicroscope. High-resolution digital images were acquired with a Nikon Digital Sight 10 camera system mounted on a Nikon Ni-E transmitted-light microscope.

For SEM analysis, two small subsamples (approximately 5 × 3 mm) containing portions of the rachis and barbs were carefully excised. These were manually polished with fine-grit sandpaper and treated with chloroform following Wang et al. (2025) to dissolve the surface matrix and expose the feather microstructures. The exposed surfaces were subsequently sputter-coated with gold and examined using a Hitachi SU5000 SEM at ECNU, operated at an accelerating voltage of 3 kV.

### Quadratic Discriminant Analysis (QDA)

To infer the coloration of specimen ECNU A0033, we quantified melanosome geometry following established protocols [24-26]. Morphometric parameters of melanosomes were measured from high-resolution SEM images using ImageJ, including length, width, and aspect ratio. Comparative reference data for melanosomes from extant avian feathers were obtained from Li et al. (2012).

All measurements were standardized prior to analysis. Quadratic discriminant analysis (QDA) was used to classify fossil melanosomes relative to extant colour categories. The model was trained on the extant dataset and subsequently applied to ECNU A0033. Classification accuracy and posterior probabilities were calculated to assess assignment confidence. Statistical analyses were performed in SPSS (version 27.0.1; IBM Corp.).

### Fluid–Structure Interaction (FSI) Analysis

A one-way fluid–structure coupling framework was used to evaluate feather deformation under aerodynamic loading, following Zhang et al. (2025). Model geometry was parameterized to approximate ECNU A0033, with a total feather length of 150 mm, barb length of 20 mm and barb orientation of 45° relative to the rachis. Two rachis configurations were constructed, each with an outer diameter of 1.7 mm, and barb cross-sections represented by ellipses with semi-major and semi-minor radii of 0.5 mm and 0.1 mm, respectively. Cylindrical rachis model: A solid cylindrical shaft with an outer diameter of 1.7 mm, representing the mechanically optimized condition seen in modern primary feathers. Open rachis model: A hollow shaft with a central cavity of 1.2 mm diameter and a cortical wall thickness of 0.25 mm, forming a semi-tubular open structure consistent with the morphology observed in A0033. In both models, barbs were represented by elliptical cross-sections (semi-major axis = 0.5 mm; semi-minor axis = 0.1 mm), and airflow was applied perpendicular to the vane plane.

### Allometric Scaling Analysis

To establish a reference framework for diameter–length scaling in modern avian primary feathers and rectrices, we analyzed primary feather datasets [27, 28] and rectrix datasets [29, 30]. All variables were log_10_-transformed. Scaling exponents were estimated using reduced major axis (RMA) regression implemented in the lmodel2 package. Ordinary least squares (OLS) regression was also performed using the lm function. 95% prediction intervals were calculated to characterize statistical variation.

In addition to the fossil specimen A0033, published measurements of Mesozoic feathers were included: *Archaeopteryx lithographica, Confuciusornis sanctus* [31], and *Caudipteryx dongi, Microraptor zhaoianus* (measured in this study on the open type rachis at 75% of feather length from the base due to poor preservation of the feather base). These fossil data were superimposed onto the modern feather scaling models to test whether they fall within the 95% prediction intervals, allowing assessment of their structural scaling patterns. All analyses were conducted in R version 4.3.1.

## Results

ECNU A0033 is a piece of amber measuring approximately 120 mm in length, 70 mm in width, and 20 mm in thickness. The preserved portion of the included feather spans approximately 105 mm, with the maximum width of the symmetric vanes reaching about 40 mm (Fig. 1A). Based on its overall proportions, the missing proximal segment is estimated to account for approximately one-quarter of its original length, suggesting that the complete feather likely exceeded 140 mm.

**Figure 1.**
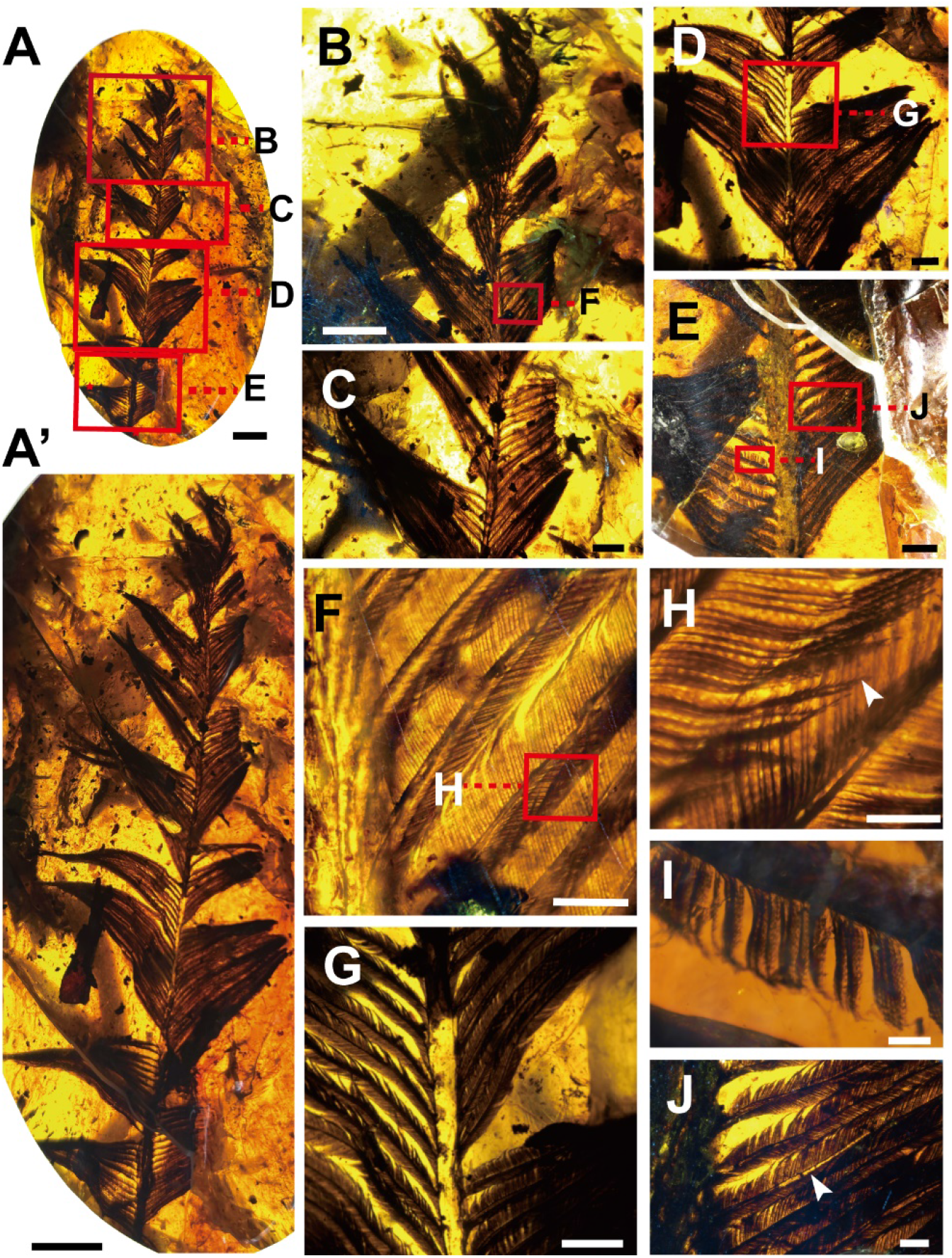
Morphology of the largest known pennaceous feather embedded in amber (ECNU A0033). **A-A’**, Photograph of the isolated pennaceous feather preserved in ECNU A0033 showing the overall shape and symmetric vanes, red boxes in **A** indicate the regions shown in **B-E**, and red stars mark the positions from where the SEM samples were excised. **B-E**, Close-up images of different portions of the feather shown in **A.** The red box in **B** marks the area shown in **F**, and the red box in **D** indicate the area shown in **G. F**, Close-up image of area highlighted in **B**, showing the closely packed barbules; **G**, Close-up images of the area highlighted in **D**, showing the rachis morphology; **H**, Close-up images of barbules showing that the proximal and distal barbules overlap directly without any evidence of hooklets; white arrowhead indicates a cilium; **I**, Close-up images showing the poorly differentiated barbule cells at the base of a barb; **J**, Close-up image showing the open barb rami (white arrowhead). **Scale bars:**10mm for **A** and **A’**; 5mm for **B**; 2.5mm for **C** and **D**; 20mm for **E**; 15mm for **G** and **J**; and 0.2mm for **H** and **I**.

The rachis reaches a maximum diameter of approximately 0.5 mm along most of its preserved length and exhibits a ventrally open configuration, as evidenced by the inverted U-shaped cross-section observed at the broken proximal end (Fig. 2A). This morphology identifies ECNU A0033 the longest known pennaceous feather preserving a ventrally open, incompletely differentiated rachis. Unlike previously described contour feathers bearing a ventrally-open rachis [19], ECNU A0033 lacks the lateral cortical expansions that would otherwise compensate for the open ventral side. Instead, the rachis shows pronounced dorsal cortical thickening (Fig. 2A). At the broken end, the dorsal cortex accounts for approximately 14% of the total rachis diameter (0.5 mm), substantially exceeding that of previously documented open-rachis specimens [18, 19].

**Figure 2.**
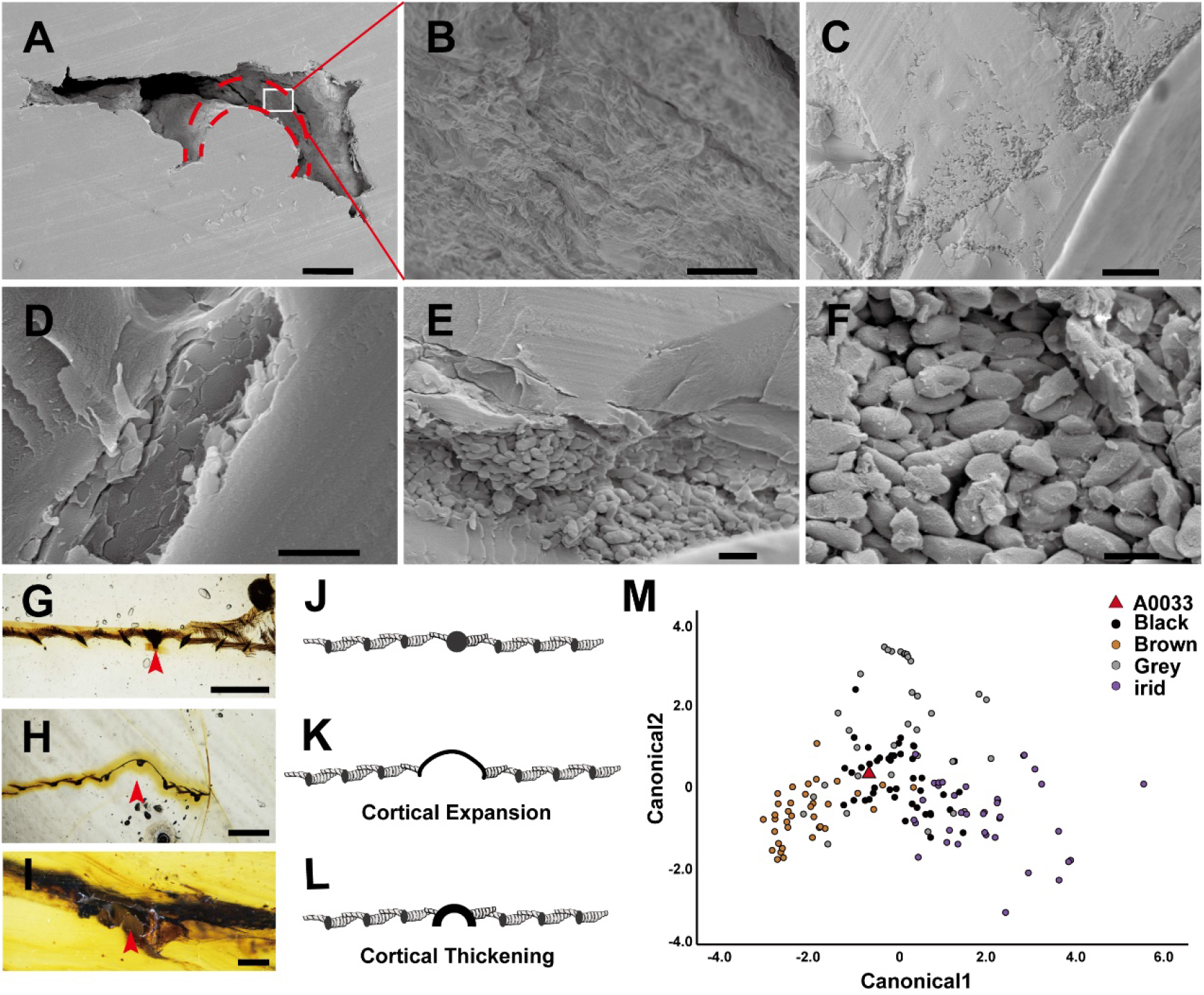
Rachial architecture and melanosomes morphology in ECNU A0033. **A-F**, SEM images of different regions of ECNU A0033. **A**, Cross-section of the feather near the preserved proximal end, showing a ventrally open rachis. The dorsal cortex was lost during SEM sampling, but its original arched outline can still be inferred (red dashed line). The white box marks the region shown in **B. B**, Carbonized matters within the cortical region, indicating the former presence of keratinized cortex. **C–F**, Melanosome preserved within barbs at different magnifications, showing their arrangement and overall morphology; **G–I**, Comparison of rachis cross-sectional architectures among amber-embedded feathers: **G**, Feather with a solid rachis (red arrowhead; CNU A0002) [19]; **H**, Feather with a ventrally open rachis and a thin dorsal cortex laterally expanded on both sides (red arrowhead; CNU A0005) [19], and **I**, ECNU A0033 showing a ventrally open rachis with a markedly thickened cortex that is not laterally expanded as in CNU A0005; **J–L**, schematic drawings of rachial cross-sections corresponding to **G–I**, highlighting structural differences. **M**, Quadratic discriminant analysis (QDA) of melanosome geometry, indicating a gray original coloration for the sampled region of ECNU A0033; **Scale bars:** 0.2mm for **A;** 10µm for **B** and **C**; 2µm for **D** and **E**; and 1µm for **F**.

The barbs are densely and regularly spaced, with partial overlap of proximal and distal barbules. Light microscopy reveals the presence of ventrally open barbs *sensu* Zhang, Tang et al. (2025). The basal segments of the barbules are relatively elongate, and although nodal cilia are observed, hooklets are absent (Fig. 1F). No evidence of barbule interlocking is observed.

SEM imaging reveals abundant elongate microbodies with a consistent aspect ratio of approximately 2:1 (Fig. 2F), consistent with the morphology of eumelanosomes typically indicative of dark pigmentation [32, 33]. Their irregular orientation is interpreted as taphonomic deformation caused by resin flow or chloroform treatment. Quadratic discriminant analysis of the melanosomes from the sampled area suggests that the original feather coloration of this region was likely gray to black (Fig. 2M, Table 2).

**Table 1.** Eigenvalues and standardized scoring coefficients for a quadratic discriminant analysis of color in extant birds and in ECNU A0033. The analysis identified six melanosome morphological and distributional properties predictive of color in extant birds, which were then used to infer the coloration of ECNU A0033.

| Canonical axis # | Eigenvalue | variation explained% | Canonical correlation | Length | Length CV | Width | Width CV | Aspect ratio | Aspect ratio skew |
| --- | --- | --- | --- | --- | --- | --- | --- | --- | --- |
| 1 | 1.065 | 72.3 | 0.718 | 2.204 | 1.38 | 1.467 | 1.05 | -1.335 | -1.341 |
| 2 | 0.367 | 24.9 | 0.518 | -1.674 | 0.708 | 2.02 | 1.828 | 1.261 | -0.441 |
| 3 | 0.04 | 2.7 | 0.169 | -0.483 | -0.229 | -0.026 | -1.156 | 1.158 | 0.918 |

**Table 2.** Color predictions for ECNU A0033 based on melanosome morphology. *P* indicates the posterior probability of group assignment; secondary predictions are shown in parentheses.

| Sample | Predicted color | *P* |
| --- | --- | --- |
| A0033 | Grey (Brown) | 0.50 (0.257) |

The FSI simulations reveal marked differences in deformation behavior between the ventrally open and cylindrical rachis configurations under identical aerodynamic loading conditions. The open-rachis model (Fig. 3A) exhibits an order-of-magnitude greater displacement than the cylindrical rachis model. In contrast, the cylindrical rachis model (Fig. 3B) shows significantly lower overall displacement and a more gradual, uniform displacement gradient along the feather length.

**Figure 3.**
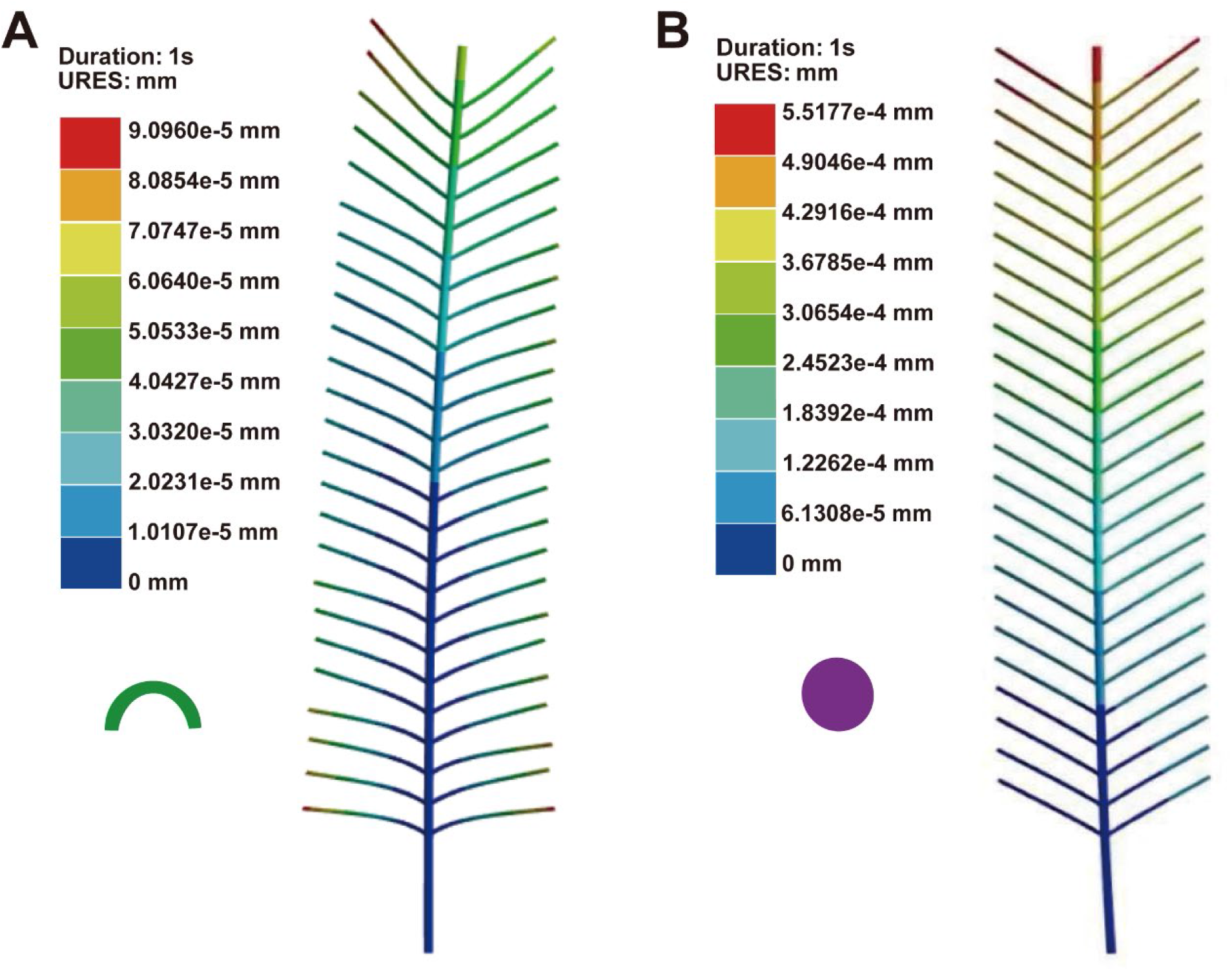
Simulated maximum displacement of feathers with ventrally open and solid rachises under a 6 m/s airflow perpendicular to the vanes. **A**, Computational model simulating a feather with a ventrally open rachis, matching the vane dimensions of ECNU A0033; **B**, Computational model simulating a feather with solid cylindrical rachis representing the mechanically optimized condition of modern pennaceous feather. Both models share identical overall geometry, barb arrangement, material properties, and airflow conditions; only the rachial structure differs. Not to scale.

Diameter–length regression models constructed from four independent datasets of extant pennaceous feathers show clear inter-dataset variation (Table 3 and Fig. 4), indicating that measurement position and sample composition exert systematic effects on the estimation of scaling parameters. Despite this variation, ECNU A0033 consistently falls below the 95% prediction interval of extant feathers in all models. This indicates that, for its inferred feather length, the rachis of ECNU A0033 is markedly thinner than expected from modern diameter– length relationships. The same result is recovered under both OLS and RMA regression methods, showing that ECNU A0033 lies outside the allometric range of both modern primary feathers and rectrices (Fig. 4).

**Table 3.** Summary of allometric regression parameters for modern avian feathers.

| Feather Type | OLS Regression Equation | RMA Regression Equation | R2 (OLS) | Sample Size (n) | OLS 95% CI (Slope) | RMA 95% CI (Slope) | Data Source |
| --- | --- | --- | --- | --- | --- | --- | --- |
| Primaries | y=−2.085+1.018x | y=−2.26+1.105x | 0.861 | 73 | [0.921, 1.115] | [1.005, 1.216] | Wang et al. (2012) |
| Primaries | y=−0.636+0.513x | y=−0.804+0.588x | 0.613 | 106 | [0.434, 0.592] | [0.501, 0.683] | Lees et al. (2017) |
| Rectrices | y=−1.958+1.043x | y=−2.214+1.17x | 0.819 | 72 | [0.926, 1.161] | [1.047, 1.311] | Dickinson et al. (2023) |
| Rectrices | y=−1.013+0.539x | y=−1.406+0.75x | 0.423 | 72 | [0.389, 0.689] | [0.559, 0.984] | Tubaro et al. (2002) |

**Figure 4.**
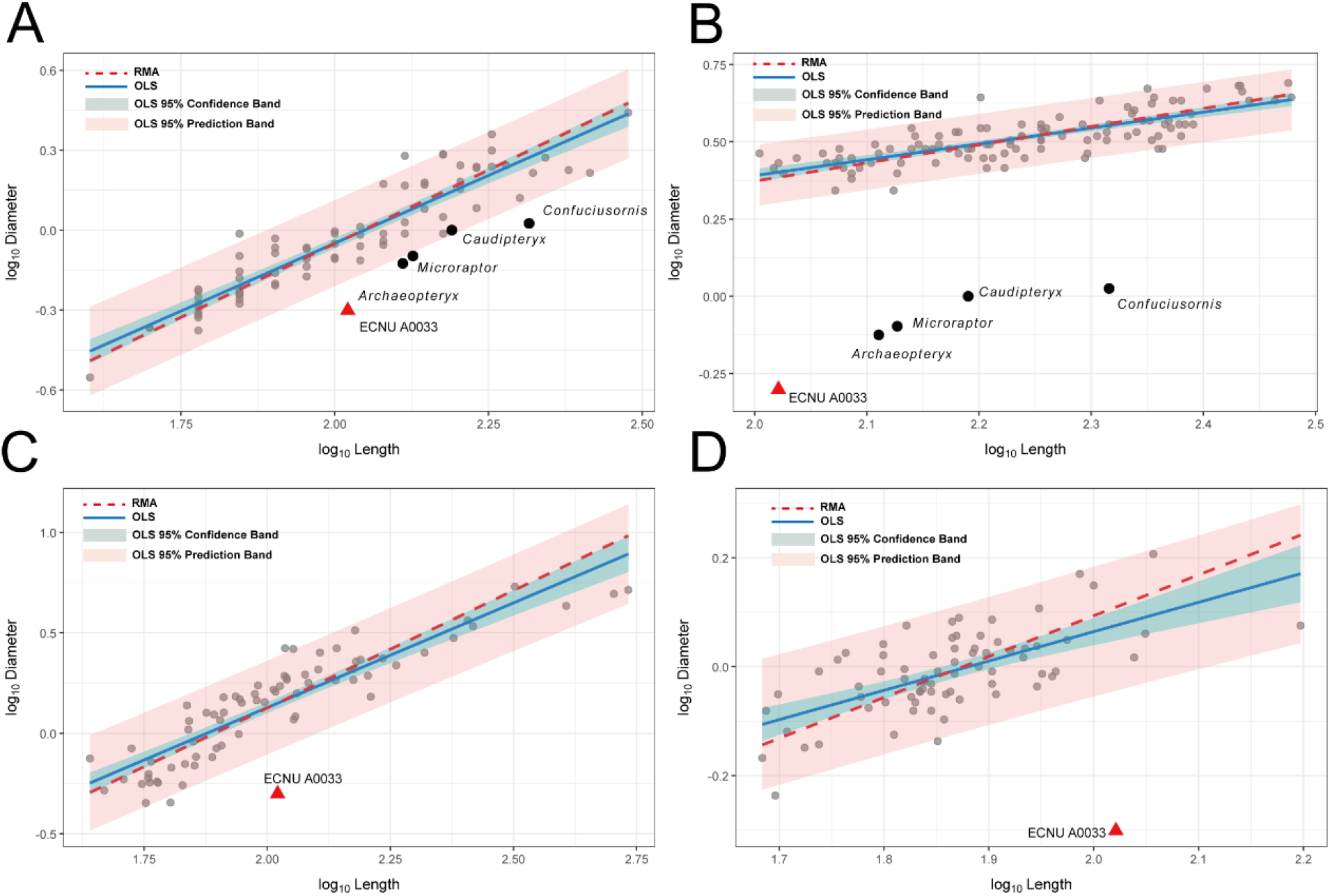
Diameter–length scaling relationships of modern avian feathers and comparison with the fossil specimen. (A) Primary feather dataset from Wang et al. (2012) (rachis diameter measured at 75% of feather length from the base); (B) Primary feather dataset from Lees et al. (2017) (rachis diameter measured at the feather base); (C) Rectrix dataset from Dickinson et al. (2023) (basal rachis diameter of central rectrices); (D) Rectrix dataset from Tubaro et al. (2002) (basal rachis diameter of central rectrices). All data (in mm) were log_10_transformed before allometric fitting: blue solid lines represent ordinary least squares (OLS) regression fits, red dashed lines represent reduced major axis (RMA) regression fits; light red transparent shading indicates the 95% prediction band of the OLS regression, light blue transparent shading indicates the 95% confidence band of the OLS regression; the red triangle denotes the amber specimen ECNU A0033, and solid black circles represent fossil data.

**Figure 5.**
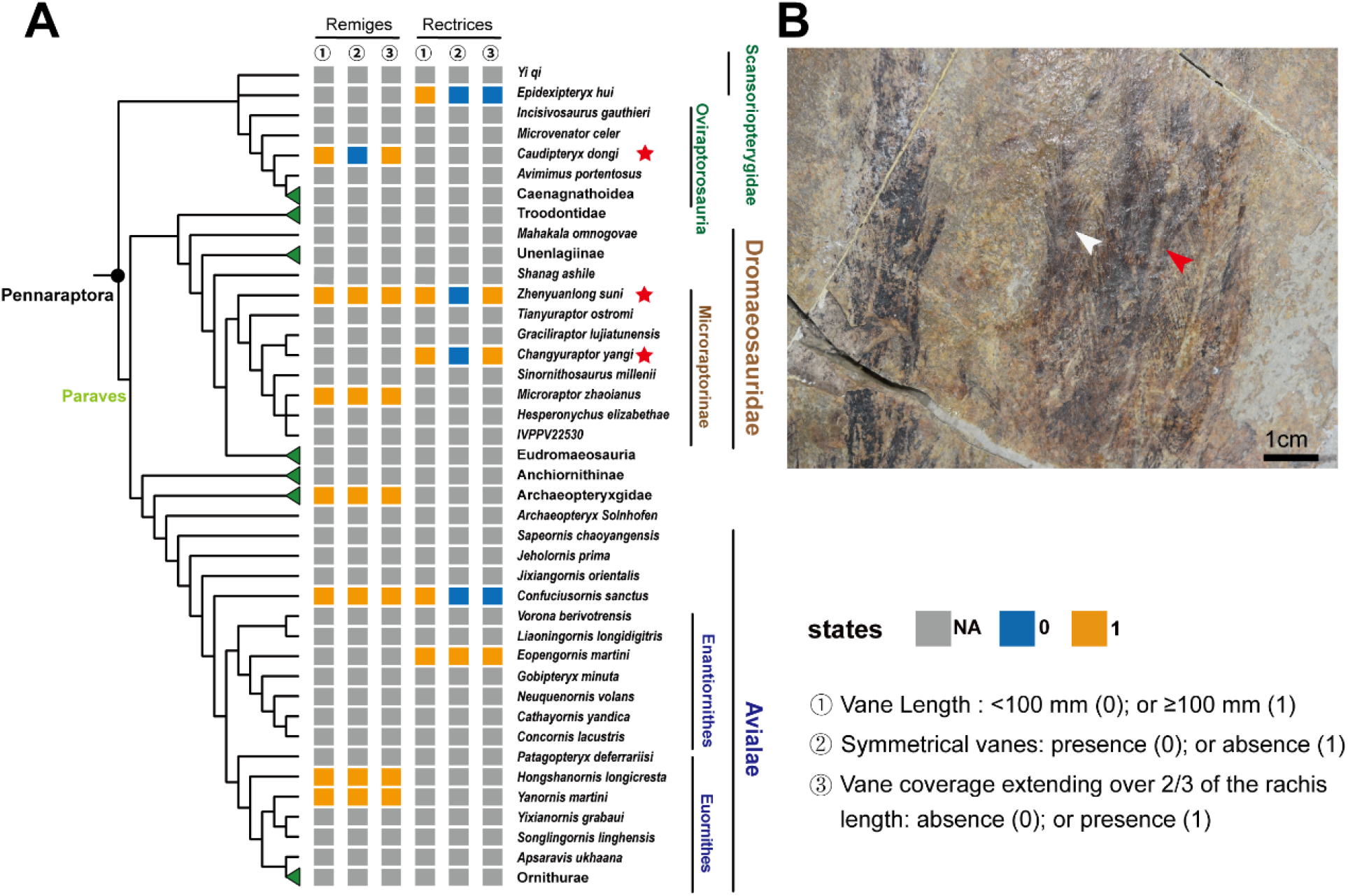
Distribution of feather characteristics observed in ECNU A0033 among pennaraptoran theropods. **A**, Distribution of three key feather traits (scored separately for remiges and rectrices) mapped onto a phylogeny of Pennaraptora. Taxa exhibiting the same combination of traits as ECNU A0033 are marked with a red star. Extant birds and fossil taxa lacking clear evidence of large pennaceous feathers were collapsed to emphasize informative nodes. The distribution indicates that the feather type of ECNU A0033 is most consistent with non-avialan pennaraptoran theropods. **B**, Primary remiges of *Caudipteryx* (IVPP V12344), showing variation in rachis structure within the series. The white arrowhead indicates a typical rachis, whereas the red arrowhead indicates a thickened rachis that, despite its ordinary appearance, may have been ventrally open, consistent with observations from a previous study [19]. This morphology represents a candidate match for the trait combination observed in ECNU A0033.

## Discussion

Morphological implications: Although the proximal portion of the vane and the calamus are not preserved in ECNU A0033, the overall shape of the preserved feather indicates that its original length exceeded 140 mm, making it by far the longest pennaceous feather discovered in amber. Its exceptional preservation enables direct assessment of feather architecture at a scale rarely captured in amber inclusions.

A particularly notable feature of ECNU A0033 is the rachis, which consists solely of a thickened dorsal cortex without a ventral cortex or internal medullary cavity. This condition has previously been reported in both small contour feathers preserved in amber and in larger compressed tail feathers (rectrices) of non-avialan and basal avialan theropods [16, 19, 34]. In contrast to previously described smaller contour feathers, where a ventrally open rachis is typically accompanied by pronounced lateral expansion of the dorsal cortex [19], ECNU A0033 shows minimal lateral expansion despite marked dorsal cortical thickening, indicating a different mode of structural reinforcement. This is notable because large feathers associated with aerodynamic function are generally expected to require greater axial stiffness than small contour feathers. In ECNU A0033, mechanical compensation appears to have been achieved primarily through dorsal cortical thickening rather than lateral expansion, suggesting that early feathers employed multiple alternative strategies to offset the absence of a ventral cortex and medullary cavity.

Importantly, ventral openness is not restricted to the rachis alone. At least in certain regions, the barbs also lack a ventral cortex, indicating that they are likewise ventrally open, a feature previously documented only in small contour feathers [18]. The co-occurrence of ventrally open rachis and barbs indicates that this feather lacked the developmental capacity to produce a fully differentiated medulla and a continuous ventral cortex [18]. This challenges a common assumption that the presence of a well-defined rachis and vane implies modern feather-like tissue differentiation [31]. Because barb microstructure is rarely preserved in compression fossils, such inferences are typically based on overall feather morphology alone. ECNU A0033 therefore provides direct fossil evidence that incomplete tissue differentiation was likely widespread in early pennaceous feathers, and that functional interpretations based solely on gross feather morphology require re-evaluation [15].

The absence of hooklets, together with the presence of nodal cilia and ventrally open barbs, indicates that ECNU A0033 lacked a fully interlocking vane. This is consistent with the interpretation that nodal cilia represent precursors to hooklets [1]. The absence of vane interlocking further implies that this feather had limited aerodynamic performance despite its large size and pennaceous appearance.

Collectively, these observations indicate that pennaceous feather evolution was asynchronous and modular, in which branching organization, elongation and vane organization preceded the acquisition of interlocking barbules and fully differentiated cortical and medullary tissues required for aerodynamic function. Although the extent to which this sequence was driven by developmental constraints remains uncertain, these results indicate that early stages of pennaceous feather evolution (including branching pattern and elongation) were not primarily shaped by aerodynamic selection, whereas later innovations such as barbule interlocking and full tissue differentiation were likely associated with increasing aerodynamic demands.

### Coloration

The dark pigmentation of ECNU A0033 is consistent with previous reports of feather coloration in non-avian theropods and early birds [25, 26], suggesting that dark or otherwise muted hues were widespread among feathered dinosaurs, including early birds. The observed dark coloration may have served functional roles such as camouflage, thermoregulation, abrasion resistance, or UV protection [35]. Given that aerodynamic performance in early pennaceous feathers was likely limited, the evolution of conspicuous or high-contrast coloration may have been constrained, as visually prominent plumage could have incurred ecological costs without providing sufficient locomotory or display advantages. ECNU A0033 therefore suggests that, even after substantial feather elongation and structural specialization, pigmentation remained relatively conservative, indicating that early pennaceous feathers were unlikely to have been primarily adapted for complex visual signaling.

### Functional implications

Previous studies indicate that an open rachis is mechanically less stable than a cylindrical one under comparable aerodynamic loading [19]. This reduced stability is also evident in ECNU A0033, despite its relatively limited rachial openness compared with previously described contour feathers. These results suggest that retention of an open rachis imposed a significant mechanical constraint on large pennaceous feathers prior to the evolution of a fully enclosed cylindrical architecture.

Although ECNU A0033 had already achieved substantial elongation and developed an overall form resembling modern flight-related feathers, the persistence of an open rachis and the absence of a fully closed vane indicate that its aerodynamic performance was likely limited. These findings demonstrate that external resemblance to modern primary feathers does not necessarily imply equivalent mechanical performance in early pennaceous feathers. The simulation results therefore reinforce the need for caution when inferring flight capability from compression fossils, especially where internal rachis architecture and barb microstructure cannot be assessed.

### Diameter–length allometry distinguishes ECNU A0033 from extant primaries and rectrices

The diameter–length proportion of ECNU A0033 falls well outside the allometric range of modern avian primary feathers and rectrices (Fig. 4). Because the proximal end is not preserved, the measured rachis diameter does not derive from the basal region typically sampled in modern feathers [28, 29]. Even under a conservative assumption that this value represents a more distal, and therefore thinner, portion of the rachis, regression models predict a total feather length of only 30∼50 mm, far shorter than the preserved length of 105 mm. Positional uncertainty therefore accounts for only part of the discrepancy. The rachis of ECNU A0033 is thus proportionally more slender than predicted by modern avian diameter–length scaling.

This result is consistent with previous observations on early feathers. Open rachises in early feather morphotypes imply incomplete tissue differentiation [19], and relatively slender rachises have also been documented in early primaries, including those of *Confuciusornis* and *Archaeopteryx* [31] as well as in other early feather morphotypes discussed by Wang et al. (2025).

Importantly, ECNU A0033 deviates from all modern reference conditions. Across four independent modern datasets, and under both OLS and RMA regression, it consistently plots below the 95% prediction interval of modern birds (Fig. 4). It therefore conforms to neither the scaling regime of extant primary feathers nor that of extant rectrices, despite superficial resemblance to both. ECNU A0033 is not simply an unusually slender analogue of a modern feather type, but occupies a region of morphospace outside the range of extant flight-related feathers.

### Taxonomic inference of the host taxon

Because ECNU A0033 falls outside the allometric range of both extant primaries and rectrices, its closest points of comparison are more likely to lie among fossil pennaceous feathers rather than among extant avian feather types. This shifts the taxonomic framework from comparison with extant feather categories to fossil clades that produced large pennaceous feathers of similar overall organization. Ventrally open rachises have been directly documented in a small number of exceptionally preserved fossil feathers and inferred in several non-avialan and avialan theropods [19]. For example, the compressed and carbonized primary feathers of *Caudipteryx* have been hypothesized to possess ventrally open rachises [19], although this interpretation remains tentative because of limited resolution and preservational quality. ECNU A0033 provides direct structural evidence supporting this hypothesis by demonstrating that feathers with apparently tubular rachises may lack a true ventral cortex and internal medulla. In such cases, the appearance of a cylindrical rachis may simply reflect the absence of obvious cortical expansion.

Building upon the latest phylogenetic framework of Pennaraptora [8, 36], we surveyed taxa bearing pennaceous feathers that meet three criteria: vane lengths exceeding 100 mm, symmetrical vanes, and vane coverage extending over more than two-thirds of the rachis length. These criteria were intended to exclude feathers with ventrally open rachises that lack vanes, as well as feathers in which vanes are restricted to the distal part of the shaft. Importantly, the presence or absence of a ventrally open rachis was not used as a selection criterion because our results indicate that many apparently cylindrical rachises may in fact prove to be ventrally open upon closer structural examination. This makes gross external morphology alone an unreliable guide to internal rachis architecture. Some pennaraptorans, such as *Similicaudipteryx* and the large-bodied *Gigantoraptor*, may also have borne very long pennaceous feathers [37, 38]. However, because such feathers have not been directly preserved in association with diagnostic skeletal material, they were not included in our survey.

This survey allows a preliminary assessment of how these feather traits are distributed across lineages and functional contexts. Pennaceous feathers exceeding 100 mm in length are not restricted to crown birds but also occurred in several non-avialan theropod clades, including microraptorines and *Caudipteryx* [39]. For instance, *Changyuraptor, Microraptor*, and *Zhenyuanlong* all possess wing or tail feathers exceeding 100 mm, with *Changyuraptor* possessing some of the longest known examples [40-42].

In extant birds, large symmetrical feathers are largely restricted to rectrices adjacent to the body midline axis. By contrast, non-avialan theropods and early birds commonly exhibit such feathers in both remiges and rectrices, suggesting a lower degree of aerodynamic specialization. When the additional criterion of symmetrical vanes extending over more than two-thirds of the rachis is applied, comparable taxa include *Caudipteryx, Zhenyuanlong, Changyuraptor*, and early avialans such as *Eopengornis* and several other taxa [43, 44]. The giant pennaceous feather preserved in ECNU A0033 likely belonged to a member of one of these clades. However, in the absence of direct skeletal association, any more specific taxonomic assignment would be unwarranted.

Rather than indicating close affinity with a particular lineage, ECNU A0033 supports the broader inference that ventrally open rachises were widespread among non-avialan theropods and basal birds, even in feathers that appear cylindrical at macroscopic resolution. The unusually thickened dorsal rachial cortex of ECNU A0033 further suggests that, in the absence of a ventral cortex and internal medulla, mechanical strength could be enhanced through alternative structural strategies. In smaller contour-like feathers, this compensation may take the form of lateral expansion of a thin dorsal cortex (Fig. 2H, K). In ECNU A0033, by contrast, it is expressed as localized dorsal thickening without substantial widening of the rachis (Fig. 2A, I, L). These contrasting strategies suggest that compensatory reinforcement of the rachis evolved prior to the establishment of a fully enclosed cylindrical architecture, although whether these differences are size-dependent remains unclear.

### Evolutionary developmental significance

Studies of extant birds have shown that feather length is not governed by a single genetic determinant, but instead reflects the interplay of at least two semi-independent developmental parameters: axial growth rate and growth duration, such that elongation can be achieved through accelerated growth, prolonged growth, or a combination of both [45].

Wu et al. (2025) showed that the FGF and IGF signaling pathways primarily promote axial elongation by enhancing cell proliferation within the feather follicle. In contrast, the Notch and Yap pathways regulate the topology and persistence of stem/progenitor cell populations, thereby delaying terminal differentiation and extending the growth phase [45]. Within this framework, the inferred length of ECNU A0033 (approaching 140 mm) indicates that at least one of these elongation-promoting mechanisms, either accelerated growth or prolonged growth duration, and quite possibly both, had already evolved.

Despite its considerable length and well-developed vanes, ECNU A0033 retains a structurally primitive rachis and barbs. This combination indicates that the developmental programs controlling feather elongation and vane elaboration were decoupled from those governing rachial cortical tissue differentiation. In other words, the mechanisms responsible for increasing feather size and shaping an elaborate vane appear to have evolved before those required for full rachial reinforcement. As the primary load-bearing element, the rachis may therefore have undergone mechanical optimization only later in feather evolution.

Developmentally, reinforcement of the rachis is achieved through coordinated epidermal differentiation and tissue remodeling. Cortical thickening results from the orderly stacking and keratinization of epidermal cells, while medullary cells undergo programmed vacuolization [46]. These processes together give rise to a lightweight, hollow, thick-walled composite beam capable of resisting aerodynamic loads. These aspects of radial differentiation are regulated by epithelial patterning pathways (notably WNT and BMP signaling), which are distinct from those controlling feather elongation [47-49]. These signaling pathways modulate the spatiotemporal expression of key keratin genes (e.g., *KRT75, FK12, KRT13A*) that guide the formation of both cortical and medullary compartments and optimize rachis mechanics for aerodynamic performance [46, 50]. This supports the idea that feather elongation and radial differentiation (i.e., structural patterning across the follicle) are regulated by distinct developmental systems.

Taken together, the combination of highly elaborate vanes and structurally primitive rachises in early feathers likely reflects a genuine asynchrony in the evolutionary timing and developmental regulation of different feather modules, rather than a taphonomic artifact [19]. This modular dissociation implies that elongation and vane elaboration could have evolved independently of, and prior to, rachial reinforcement. ECNU A0033 therefore supports a stepwise developmental sequence in early pennaceous feather evolution (Fig. 6), in which elongation and vane organization preceded the acquisition of fully differentiated cortical– medullary architecture and aerodynamic refinement.

**Figure 6.**
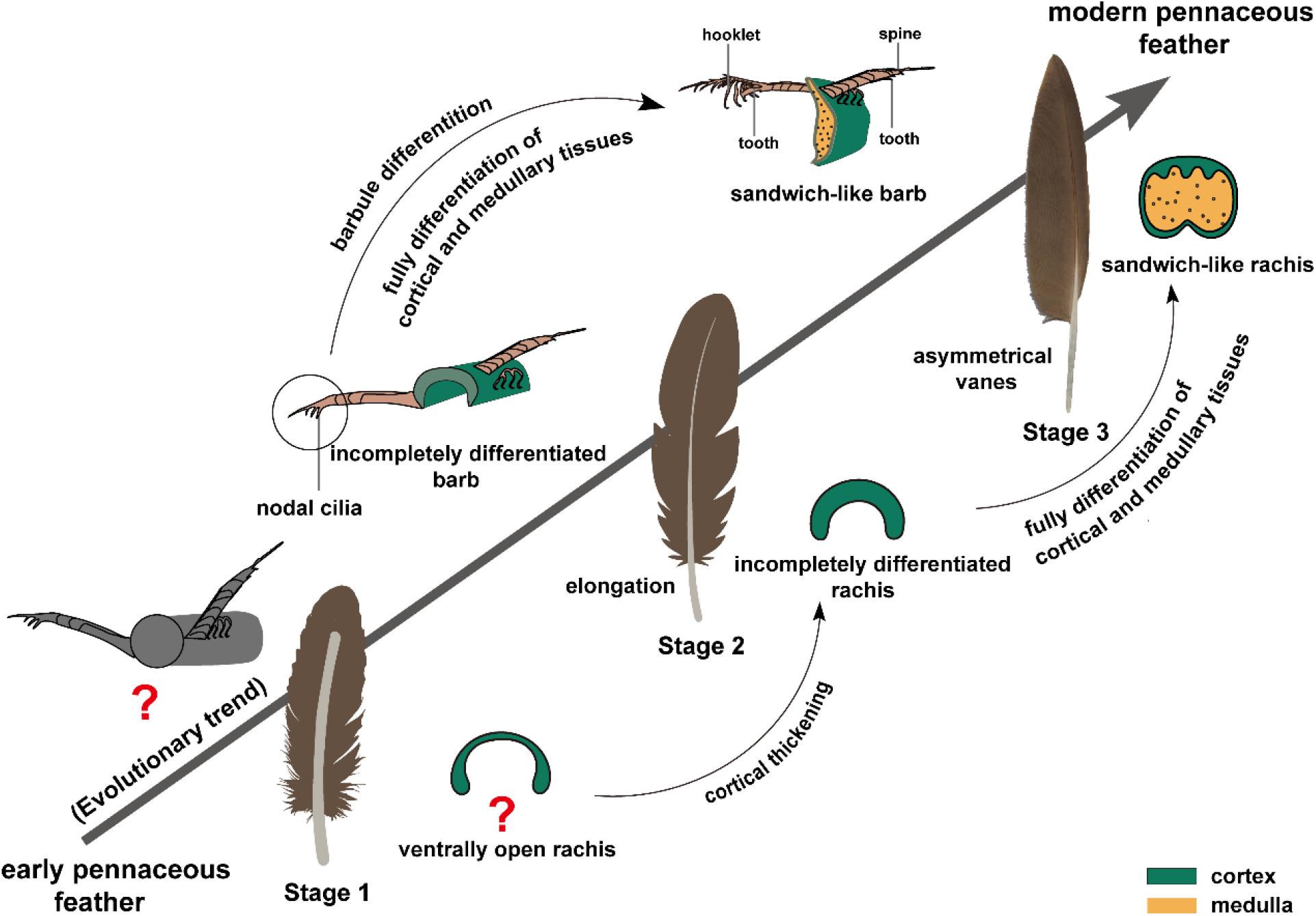
Stepwise and modular evolution of pennaceous feathers. This schematic illustrates the proposed evolutionary sequence from early pennaceous feathers to modern forms, highlighting the decoupled development of branching architecture, elongation and tissue differentiation. Stage 1: Hypothesized early pennaceous feather with poorly differentiated barbs and rachis, lacking clear cortical–medullary differentiation. Stage 2 (ECNU A0033): Transitional stage with a ventrally open rachis and rami, showing dorsal cortical thickening but reduced lateral expansion. Barbs bear nodal cilia but lack hooklets and interlocking barbules, resulting in an incompletely closed vane. Stage 3: Fully differentiated feather with a closed rachis and barbs exhibiting a sandwich-like cortical–medullary structure, and well-developed hooklets enabling barbule interlocking. Arrows indicate inferred evolutionary processes, including cortical thickening, complete cortical–medullary differentiation, and the acquisition of barbule interlocking.

## Conclusion

ECNU A0033 provides direct fossil evidence that pennaceous feather evolution proceeded in a stepwise and modular manner. Its morphology shows that major components of pennaceous feather construction did not arise simultaneously: substantial elongation and a well-developed symmetrical vane were already established, whereas interlocking vane closure, cortical-medullary organization, and full rachial mechanical refinement remained incomplete. This decoupled pattern indicates that feathers could achieve a superficially advanced overall appearance prior to the establishment of the tissue differentiation required for effective aerodynamic function. These results further suggest that early evolution of pennaceous feather was not primarily governed by aerodynamic pressure, whereas aerodynamic demands became increasingly important with the later acquisition of interlocking barbules and fully differentiated tissue architecture.

## Acknowledgments

We thank Z.G. (ECNU) and the East China Normal University Multi-Functional Platform for Innovation (004) for SEM imaging support. S.W. was supported by the Zijiang Program for Talented Scholars at East China Normal University, and N.J. was supported by the Shanghai Sci-Tech Co-research Program (25HB2712800).

## Author contributions

S.W. conceived the project; Y. Z. and J.T. performed the experiments and analyzed the data; Y.Z. and S.W. wrote the paper; N.J. contributed to manuscript revision. All authors read and approved the final version of the manuscript.

